## Supplementary Figures and Tables for "Transcriptional profiling of Hutchinson-Gilford Progeria syndrome fibroblasts reveals deficits in mesenchymal stem cell commitment to differentiation related to early events in endochondral ossification"

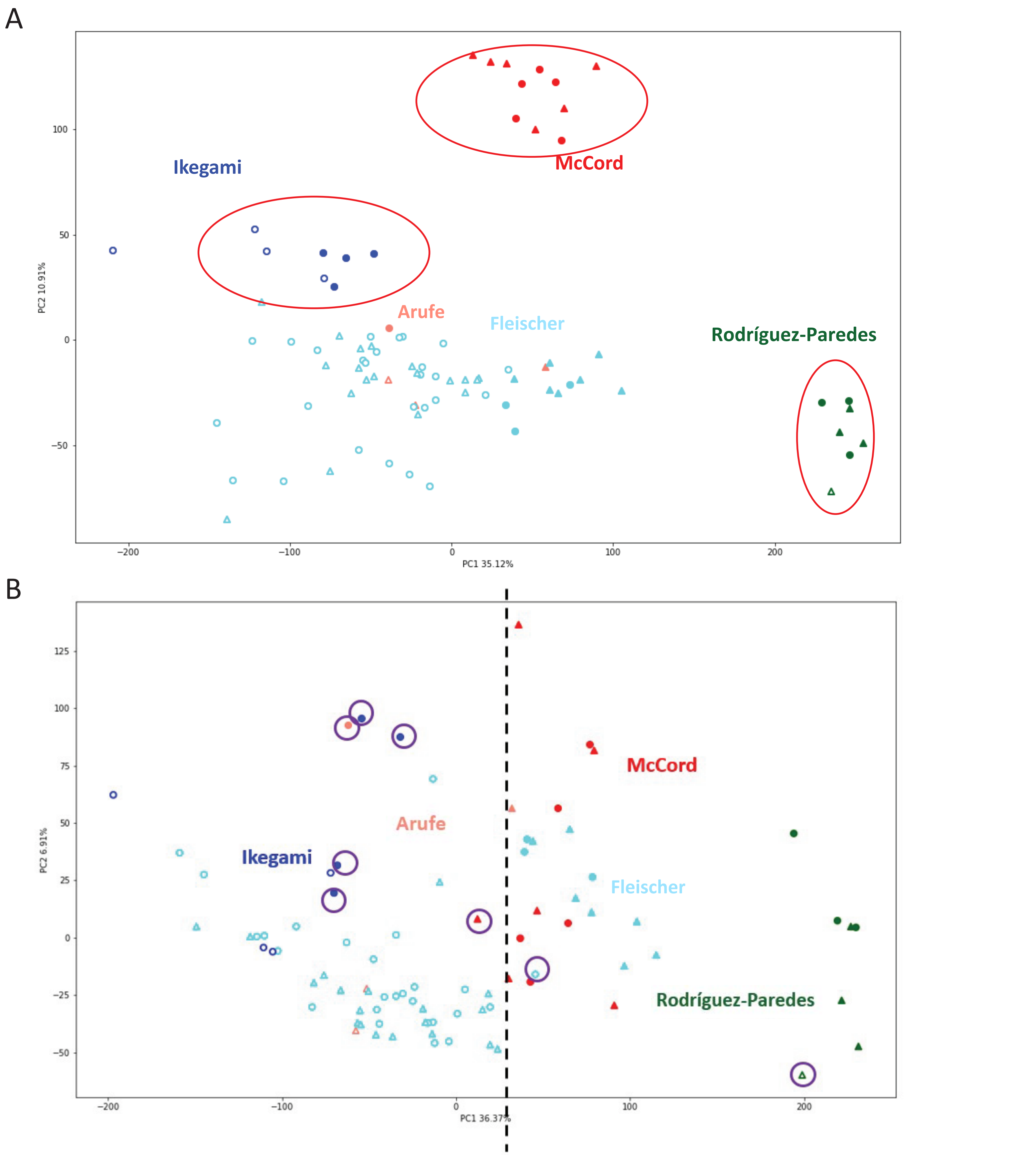


**Supplementary Figure 1. Batch clustering correction.** A) Principal component analysis (PCA) of the datasets used in this study before batch correction. Samples segregate primarily by lab of origin (red circles). Solid color markers: Progeria, hollow markers: normal controls, triangles: female, circles: male. B) PCA after batch correction. Samples primarily segregate by progeria/control phenotype. Dotted line denotes an arbitrary margin between cohorts. Solid color markers: Progeria, hollow markers: normal controls, triangles: female, circles: male. Purple circles denote patients that are mismatched from the rest of their respective groups.

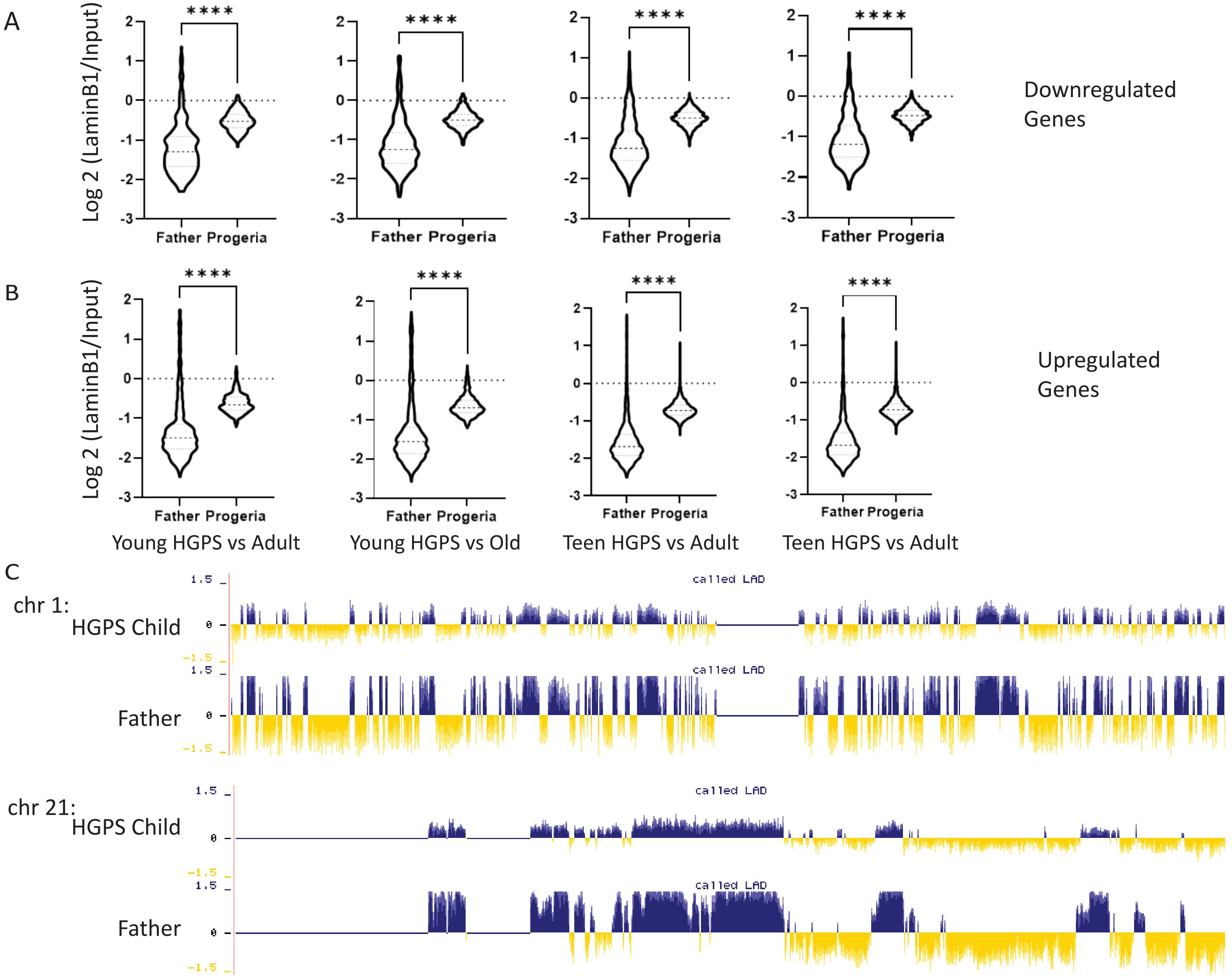


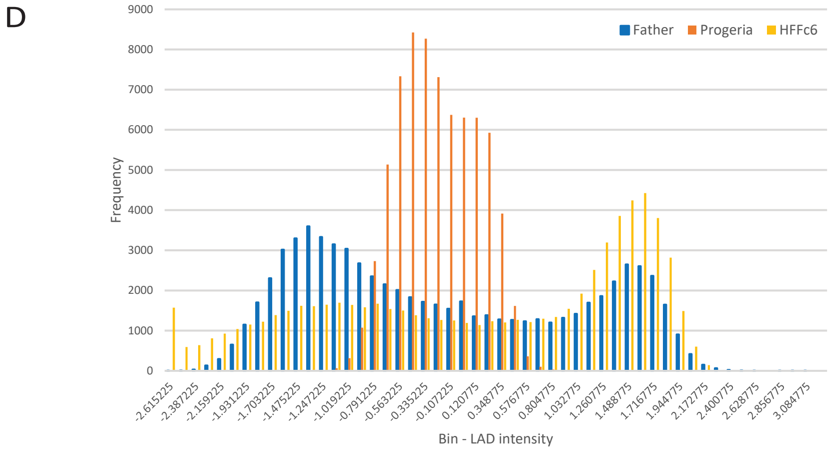


**Supplementary Figure 2.** A) The genomic regions for down-regulated genes identified in our age comparisons were compiled with the LAD identity found in a father-HGPS child fibroblasts (HGADFN168 and HGADFN167, respectively). Comparisons are presented as follows, from left to right: Young HGPS (0-7 years old) compared to middle-aged controls, young HGPS patients compared to old controls, teenaged HGPS compared to middle-aged controls and teenaged HGPS compared to old controls. Differences between al groups are significant (Kruskal-Wallis $p<0.001$) Violin plots for the global distribution of values; median denoted by thick dashed line, 25th and 75th percentiles highlighted as thin dashed lines.
B) The genomic regions for up regulated genes identified in our age comparisons were compiled with the LAD identity found in Father or HGPS child fibroblasts (HGADFN168 and HGADFN167, respectively). Comparisons are presented as follows, from left to right: Young HGPS (0-7 years old) compared to middle-aged controls, young HGPS patients compared to old controls, teenaged HGPS compared to middle-aged controls and teenaged HGPS compared to old controls. Differences between al groups are significant (Kruskal-Wallis $p<0.001$) Violin plots for the global distribution of values; median denoted by thick dashed line, 25th and 75th percentiles highlighted as thin dashed lines
C) LAD distribution along chromosome 1 and chr 21. LAD identity remains consistent among samples, with small regional changes in LAD definition in specific areas (highlighted- rectangle). The greatest difference among samples is the strength in LAD definition, as characterized by smaller positive and negative values D) The genome-wide distribution of LADs, as characterized by DamID-seq shows that normal fibroblast lines HGADFN168 and HFFc6 (human foreskin fibroblast), show a characteristic bimodal distribution in LAD intensity around positive (LAD) and negative (non-LAD) values. In contrast, LAD values for the HGPS cell line HGADFN168 distribute normally, around zero.

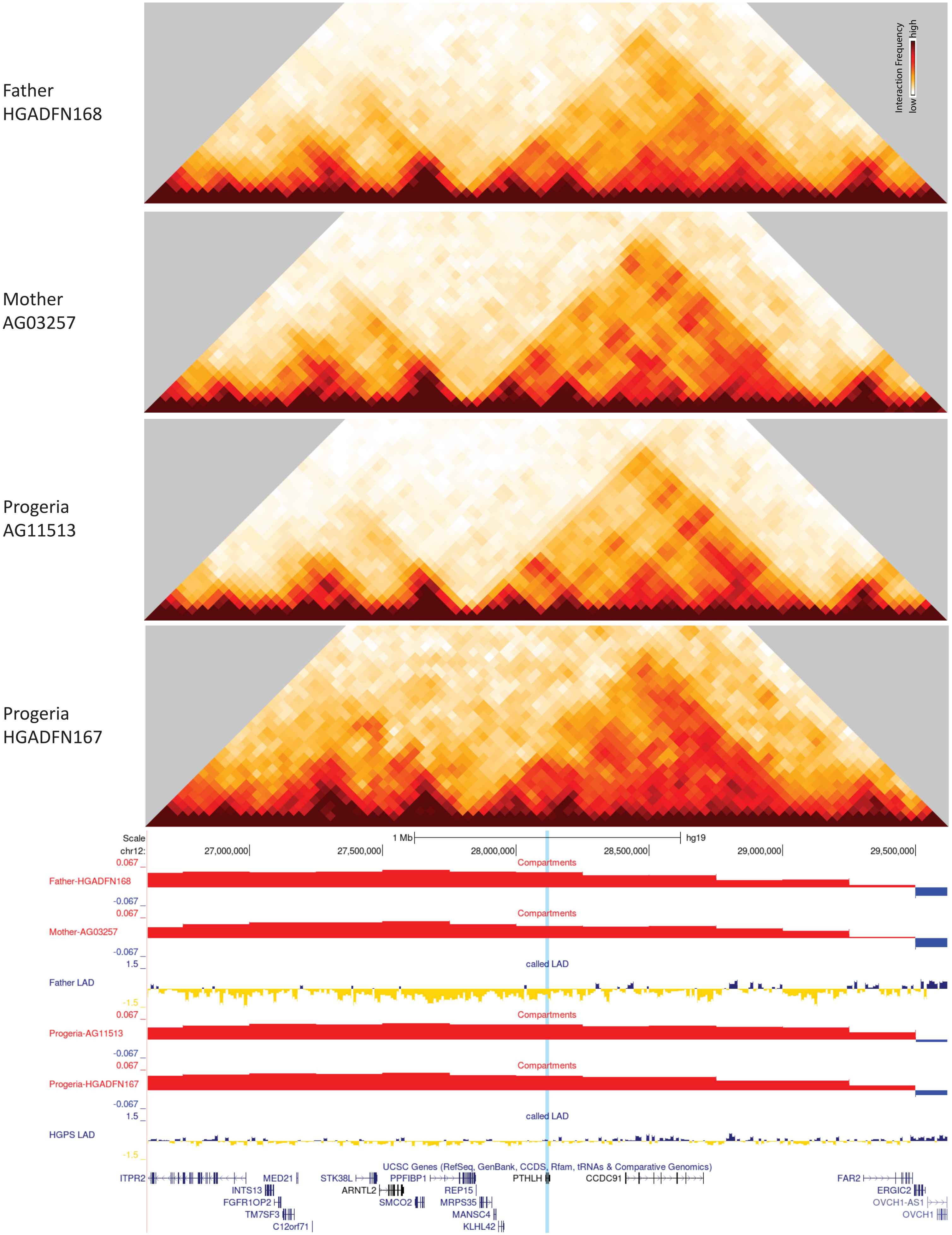


**Supplementary Figure 3.** 40-kb resolution heatmaps for the parental and HGPS cell lines, around the PTHLH gene (highlight: blue), aligned to their associated compartment and LAD tracks. The gene localizes to the A compartment in both HGPS and control cell lines. A called LAD domain is present in HGPS cells.

Supplementary Table 1

| Progeria Patient | Age | Sex | RNA-seq approach | Lab/First Author | GEO series |
|---|---|---|---|---|---|
| HGADFN155 | 1 yr 2 mo | F | polyA (TruSeq RNA Sample Preparation v2) | Rodríguez-Paredes | GSE150137 |
| HGADFN188 | 2 yr 3 mo | F | polyA (TruSeq Stranded mRNA) | Fleischer | GSE113957 |
| HGADFN188 | 2 yr 3 mo | F | polyA (TruSeq RNA Sample Preparation v2 ) | Rodríguez-Paredes | GSE150137 |
| HGADFN367 | 3 yr | F | polyA (TruSeq Stranded mRNA) | Fleischer | GSE113957 |
| HGADFN127 | 3 yr 9 mo | F | polyA (TruSeq Stranded mRNA) | Fleischer | GSE113957 |
| HGADFN164 | 4 yr 8 mo | F | polyA (TruSeq Stranded mRNA) | Fleischer | GSE113957 |
| HGADFN164 | 4 yr 8 mo | F | polyA (TruSeq RNA Sample Preparation v2) | Rodríguez-Paredes | GSE150137 |
| HGADFN122 | 5 yr | F | polyA (TruSeq Stranded mRNA) | Fleischer | GSE113957 |
| HGADFN178 | 6 yr 11 mo | F | polyA (TruSeq Stranded mRNA) | Fleischer | GSE113957 |
| AG11513 | 8 yr | F | polyA (TruSeq Stranded mRNA) | Fleischer | GSE113957 |
| HGADFN167 | 8 yr 5 mo | M | polyA (TruSeq Stranded mRNA) | Fleischer | GSE113957 |
| HGADFN167 | 8 yr 5 mo | M | polyA (TruSeq RNA Sample Preparation v2) | Rodríguez-Paredes | GSE150137 |
| HGADFN167 | 8 yr 5 mo | M | polyA (NEBNext Ultra DNA Library Prep Kit) | Ikegami | GSE113343 |
| HGADFN167-2 | 8 yr 5 mo | M | polyA (NEBNext Ultra DNA Library Prep Kit) | Ikegami | GSE113343 |
| HGADFN169 | 8 yr 6 mo | M | polyA (TruSeq Stranded mRNA) | Fleischer | GSE113957 |
| HGADFN169 | 8 yr 6 mo | M | polyA (TruSeq RNA Sample Preparation v2) | Rodríguez-Paredes | GSE150137 |
| HGADFN143 | 8 yr 10 mo | M | polyA (TruSeq Stranded mRNA) | Fleischer | GSE113957 |
| HGADFN143 | 8 yr 10 mo | M | polyA (TruSeq RNA Sample Preparation v2) | Rodríguez-Paredes | GSE150137 |
| AG03199 | 10 yr | F | polyA (Illumina SureSelect Strand Specific RNA library Prep) | Arufe | GSE113648 |
| AG03513 | 13 yr | M | polyA (Illumina SureSelect Strand Specific RNA library Prep) | Arufe | GSE113648 |
| AG11498 | 14 yr | M | polyA (NEBNext Ultra DNA Library Prep Kit) | Ikegami | GSE113343 |
| AG11498-2 | 14 yr | M | polyA (NEBNext Ultra DNA Library Prep Kit) | Ikegami | GSE113343 |

Supplementary Table 2

| Parent | Age | Sex | RNA-seq approach | Lab/Author | GEO series |
|---|---|---|---|---|---|
| AG03257 | 35 | F | polyA (Illumina SureSelect Strand Specific RNA library Prep) | Arufe | GSE113648 |
| AG03512 | 41 | F | polyA (Illumina SureSelect Strand Specific RNA library Prep) | Arufe | GSE113648 |
| **Middle age control** | | | | | |
| AG07124 | 26 | F | polyA (TruSeq Stranded mRNA) | Fleischer | GSE113957 |
| AG09599 | 30 | F | polyA (TruSeq Stranded mRNA) | Fleischer | GSE113957 |
| GM04503 | 31 | F | polyA (TruSeq Stranded mRNA) | Fleischer | GSE113957 |
| GM04504 | 31 | F | polyA (TruSeq Stranded mRNA) | Fleischer | GSE113957 |
| GM00043 | 32 | F | polyA (TruSeq Stranded mRNA) | Fleischer | GSE113957 |
| GM01650 | 37 | F | polyA (TruSeq Stranded mRNA) | Fleischer | GSE113957 |
| AG16358 | 41 | F | polyA (TruSeq Stranded mRNA) | Fleischer | GSE113957 |
| GM00495 | 29 | M | polyA (TruSeq Stranded mRNA) | Fleischer | GSE113957 |
| AG07478 | 29 | M | polyA (TruSeq Stranded mRNA) | Fleischer | GSE113957 |
| AG09605 | 30 | M | polyA (TruSeq Stranded mRNA) | Fleischer | GSE113957 |
| GM01717 | 39 | M | polyA (TruSeq Stranded mRNA) | Fleischer | GSE113957 |
| AG13967 | 41 | M | polyA (TruSeq Stranded mRNA) | Fleischer | GSE113957 |
| AG04063 | 43 | M | polyA (TruSeq Stranded mRNA) | Fleischer | GSE113957 |
| AG04054 | 29 | M | polyA (TruSeq Stranded mRNA) | Fleischer | GSE113957 |
| **Old Adult Control** | | | | | |
| GM01706 | 82 | F | polyA (TruSeq Stranded mRNA) | Fleischer | GSE113957 |
| AG04059 | 96 | M | polyA (TruSeq Stranded mRNA) | Fleischer | GSE113957 |
| AG09602 | 92 | F | polyA (TruSeq Stranded mRNA) | Fleischer | GSE113957 |
| AG08433 | 94 | M | polyA (TruSeq Stranded mRNA) | Fleischer | GSE113957 |
| AG07725 | 91 | M | polyA (TruSeq Stranded mRNA) | Fleischer | GSE113957 |
| AG05247 | 87 | F | polyA (TruSeq Stranded mRNA) | Fleischer | GSE113957 |
| AG04064 | 92 | M | polyA (TruSeq Stranded mRNA) | Fleischer | GSE113957 |
| GM03525 | 80 | F | polyA (TruSeq Stranded mRNA) | Fleischer | GSE113957 |
| AG04662 | 87 | M | polyA (TruSeq Stranded mRNA) | Fleischer | GSE113957 |
| AG11744 | 84 | F | polyA (TruSeq Stranded mRNA) | Fleischer | GSE113957 |
| AG12788 | 90 | M | polyA (TruSeq Stranded mRNA) | Fleischer | GSE113957 |
| AG11725 | 84 | F | polyA (TruSeq Stranded mRNA) | Fleischer | GSE113957 |
| AG13129 | 89 | M | polyA (TruSeq Stranded mRNA) | Fleischer | GSE113957 |
| AG04386 | 83 | M | polyA (TruSeq Stranded mRNA) | Fleischer | GSE113957 |
| AG05274 | 84 | M | polyA (TruSeq Stranded mRNA) | Fleischer | GSE113957 |

### Supplementary Table 3

| Healthy Child Control | Age | Sex | RNA-seq approach | Lab/First Author | GEO series |
|---|---|---|---|---|---|
| AG08498 | 1 | M | polyA (TruSeq Stranded mRNA) | Fleischer | GSE113957 |
| AG16409 | 12 | M | polyA (TruSeq Stranded mRNA) | Fleischer | GSE113957 |
| GM00969 | 2 | F | polyA (TruSeq Stranded mRNA) | Fleischer | GSE113957 |
| GM05565 | 3 | M | polyA (TruSeq Stranded mRNA) | Fleischer | GSE113957 |
| GM00498 | 3 | M | polyA (TruSeq Stranded mRNA) | Fleischer | GSE113957 |
| GM00038 | 9 | F | polyA (TruSeq Stranded mRNA) | Fleischer | GSE113957 |
| GM01652 | 11 | F | polyA (TruSeq Stranded mRNA) | Fleischer | GSE113957 |
| GM07753 | 17 | M | polyA (TruSeq Stranded mRNA) | Fleischer | GSE113957 |
| GM00499 | 8 | M | polyA (TruSeq Stranded mRNA) | Fleischer | GSE113957 |
| GM07532 | 16 | F | polyA (TruSeq Stranded mRNA) | Fleischer | GSE113957 |
| GM05400 | 6 | M | polyA (TruSeq Stranded mRNA) | Fleischer | GSE113957 |
| GM00969 | 2 | F | polyA (TruSeq RNA Sample Preparation v2 protocol) | Rodríguez-Paredes | GSE150137 |
| GM08398 | 8 | M | polyA (NEBNext Ultra DNA Library Prep Kit) | Ikegami | GSE113343 |
| GM07492 | 17 | M | polyA (NEBNext Ultra DNA Library Prep Kit) | Ikegami | GSE113343 |
| GM08398-2 | 8 | M | polyA (NEBNext Ultra DNA Library Prep Kit) | Ikegami | GSE113343 |
| GM07492-2 | 17 | M | polyA (NEBNext Ultra DNA Library Prep Kit) | Ikegami | GSE113343 |
| GM08398 | 8 | M | polyA (TruSeq Stranded mRNA) | Fleischer | GSE113957 |
| GM07492 | 17 | M | polyA (TruSeq Stranded mRNA) | Fleischer | GSE113957 |
| GM01582 | 11 | F | polyA (TruSeq Stranded mRNA) | Fleischer | GSE113957 |
| GM08399 | 19 | F | polyA (TruSeq Stranded mRNA) | Fleischer | GSE113957 |
| GM00409 | 7 | M | polyA (TruSeq Stranded mRNA) | Fleischer | GSE113957 |

**Supplementary Table 4. Hi-C Mapping Statistics**

| Sample Name | Enzyme | Genotype | Gender, Age | Raw Reads | Both Sides Mapped | % Mapped | %Dangling Ends | Valid Pairs | Unique Valid Pairs | %Cis |
|---|---|---|---|---|---|---|---|---|---|---|
| Progeria-167-DpnII-P19-R1 | DpnII | HGPS | Male, 8Y | 364,961,495 | 220,942,665 | 60.54 | 3.05 | 208,963,277 | 152,974,533 | 54.69 |
| Progeria-168-DpnII-P16-R1 | DpnII | WT | Male, 40Y | 358,703,556 | 220,880,406 | 61.58 | 3.48 | 212,152,596 | 148,022,234 | 54.98 |
| Progeria-AG03257-P7-R1 | DpnII | WT | Female, 35Y | 274,364,560 | 177,877,476 | 64.83 | 0.95 | 174,730,268 | 132,694,872 | 49.65 |
| Progeria-AG03257-P7-R3 | DpnII | WT | Female, 35Y | 127,656,906 | 73,190,497 | 57.33 | 2.26 | 70,541,316 | 46,463,429 | 83.56 |
| Progeria-AG11513-P7-R1 | DpnII | HGPS | Female, 8Y | 251,483,696 | 154,772,051 | 61.54 | 0.46 | 153,219,873 | 116,077,671 | 48.77 |
| Progeria-AG11513-P7-R2 | DpnII | HGPS | Female, 8Y | 251,776,055 | 153,160,225 | 60.83 | 0.75 | 151,305,617 | 114,631,955 | 50.22 |
| Progeria-HGADFN167-P12-R1 | HindIII | HGPS | Male, 8Y | 165,040,682 | 116,708,094 | 70.71 | 24.91 | 86,987,628 | 81,047,934 | 67.32 |
| Progeria-HGADFN167-P19-R1 | HindIII | HGPS | Male, 8Y | 202,090,523 | 136,172,100 | 67.38 | 14.12 | 116,222,620 | 95,537,352 | 82.72 |
| Progeria-HGADFN168-P12-R1 | HindIII | WT | Male, 40Y | 194,414,002 | 136,139,915 | 70.03 | 16.61 | 112,990,317 | 77,889,695 | 81.23 |
| Progeria-HGADFN168-P27-R1 | HindIII | WT | Male, 40Y | 157,637,031 | 110,558,519 | 70.13 | 15.06 | 93,504,712 | 86,842,660 | 67.32 |
